# Identification of major quantitative trait loci for parthenocarpic ability in East Asian melon

**DOI:** 10.1101/2023.02.16.528896

**Authors:** Yosuke Yoshioka, Akito Nashiki, Ahmad Zaelani, Sachiko Isobe, Kenta Shirasawa, Koichiro Shimomura, Akio Ohyama

**Affiliations:** Institute of Life and Environmental Sciences, University of Tsukuba, Tsukuba, Ibaraki 305-8572, Japan; Graduate School of Science and Technology, University of Tsukuba, Tsukuba, Ibaraki 305-8572, Japan; Kazusa DNA Research Institute, Kisarazu, Chiba 292-0818, Japan; Ano Vegetable Research Station, Institute of Vegetable and Floriculture Science, National Agriculture and Food Research Organization, Tsu, Mie 514-2392, Japan; Institute of Vegetable and Floriculture Science, National Agriculture and Food Research Organization, Tsukuba, Ibaraki 305-8519, Japan

**Keywords:** *Cucumis melo*, CAPS marker, dd-RAD-seq, marker-assisted selection

## Abstract

Natural (genetic) parthenocarpy contributes to fruit yield and quality under unfavorable environmental conditions where there are no effective pollinators or fertile pollen grains. Several old melon cultivars and weedy melon in East Asia are known to have strong parthenocarpic ability, but there has been little progress in intentionally introducing this ability into current F_1_ hybrid cultivars. Here, we examined its inheritance and confirmed the selection accuracy of DNA markers linked to it. We conducted QTL analysis using three F_2_ populations derived from crosses between a non-parthenocarpic cultivar and three parthenocarpic accessions, and detected two major QTLs on chromosomes 2 (*par2.1*) and 3 (*par3.1*). The parthenocarpic parent allele at both QTLs enhanced parthenocarpic ability. Phenotypic segregation was well explained by *par2.1* and *par3.1* in two F_2_ populations derived from Japanese weedy melon and an old Korean cultivar and by *par3.1* in one from an old Japanese cultivar. This difference suggests that the effects of *par2.1* and *par 3.1* depend on genetic background. Both QTL regions contain several phytohormone-related genes, so we randomly selected SNPs in auxin- and ethylene-related genes to confirm the accuracy of selection for parthenocarpic ability. These SNP markers proved sufficient, though not perfect, to select plants with strong parthenocarpic ability. These results provide new insights into the molecular mechanisms of parthenocarpic ability in melon and will contribute to the development of new cultivars with high parthenocarpic ability.

**Key message:** Several oriental melons have strong parthenocarpic ability controlled by one or two loci. DNA markers linked to these loci can select individuals with this ability.

## Introduction

Natural (genetic) parthenocarpy, an ability to develop fruits without pollination or any artificial stimuli, is a desirable trait in many fruit and vegetable crops, as it can lead to stable fruit production even in environments with few or no effective pollinators. Many farmers rely on insects (such as honeybees and bumblebees) to keep fruit production costs low, but the numbers of insect pollinators in nature have significantly decreased in recent years, and large-scale losses of managed honeybees have occurred (Potts et al. 2010; LeBuhn and Vargas Luna 2021; Zattara and Aizen 2021). In addition, low temperature during winter and early spring reduces not only pollinator activity but also the amount of fertile pollen available. These factors reduce yields and fruit quality and lengthen the cropping period. The cultivation of parthenocarpic cultivars is labor-saving because no artificial crossing or chemical treatment for inducing parthenocarpy is required. Therefore, the development and use of parthenocarpic cultivars is a promising way to sustain fruit production in unfavorable environments. Within cucurbit crops, parthenocarpy is widespread in cucumber (*Cucumis sativus* L.), and almost all current F_1_ hybrid cultivars set fruit parthenocarpically. Parthenocarpy has also been reported in squash (*Cucurbita pepo* L. and *C. moschata* Duch.) germplasms (Robinson 1993; Robinson and Reiners 1999; Takisawa et al. 2021), and several parthenocarpic cultivars have been released. Within melon (*Cucumis melo* L.), a weedy melon in the horticultural group *agrestis* and several old oriental melon cultivars in the *conomon* and *makuwa* groups have parthenocarpic ability (Yoshioka et al. 2018). However, there has been little progress in intentionally introducing the parthenocarpic ability of these cultivars/lines into current F_1_ hybrid cultivars. To efficiently introduce this trait in practical breeding, it is of great importance to understand its inheritance and to develop techniques such as marker-assisted selection.

The genetic mechanism of parthenocarpy in cucurbits is not fully understood, although some highly parthenocarpic cultivars have been developed. Recent studies suggest that parthenocarpy in cucumber is inherited in a quantitative manner and is regulated by multiple genes with different effects. Sun et al. (2006) identified quantitative trait loci (QTLs) associated with parthenocarpy, distributed across four linkage groups, in US processing cucumber lines. Lietzow et al. (2016) detected seven QTLs for parthenocarpic ability in US processing cucumber lines genetically similar to those of Sun et al. (2006), and the effects of four of those QTLs were stable in multiple environments. Wu et al. (2016) identified a major-effect QTL on chromosome (Chr.) 2 and six minor-effect QTLs in a population derived from a cross between parthenocarpic and non-parthenocarpic inbred lines derived from different two European greenhouse cultivars. Among these QTLs, only the major-effect QTL on Chr. 2 had a location largely consistent with the QTLs detected in Lietzow et al. (2016). More recently, Gou et al. (2022) performed genome-wide association study analysis of parthenocarpic ability using diverse cucumber accessions, including North and South China cucumbers, and detected several QTLs on Chrs. 3 and 6, both in common with the above studies and different ones on other chromosomes. The discrepancies may reflect not only the different lines, but also the environmental conditions or phenotyping methods used in these studies.

Parthenocarpic fruits are usually induced by exogenous application of plant growth regulators such as cytokinins, auxins, and gibberellins (Fu et al., 2008; Su et al. 2021). Yin et al. (2006) showed that enhanced expression of an auxin-synthesizing gene significantly enhanced parthenocarpic ability in cucumber. Other phytohormones and their related genes are also considered to be involved in natural parthenocarpy in cucumber (Li et al. 2014; Su et al. 2021). In addition, signal transduction pathways that do not involve phytohormones may act in some cultivars/lines (Li et al. 2017). Squash germplasms with parthenocarpic ability were reported many years ago (Robinson 1993; Robinson and Reiners 1999), and inheritance of parthenocarpy in squash has been investigated (Menezes et al. 2005; Nogueira et al. 2011). Recently, Martínez et al. (2014) reported promising zucchini accessions and associated their parthenocarpy with downregulation of ethylene production in unpollinated fruits. Pomares-Viciana et al. (2017, 2019) analyzed differences in fruit transcriptomes between parthenocarpic and non-parthenocarpic zucchini cultivars, and concluded that phytohormones play an important role: i.e., activation by auxins and gibberellins and inhibition by ethylene, in both pollinated and parthenocarpic fruits.

The inheritance of parthenocarpic ability in the cultivars and weedy melon reported by Yoshioka et al. (2018) might be similar to that in cucumber and zucchini. However, genetic analysis of parthenocarpic ability in these cultivars/lines has not progressed. Understanding the inheritance of parthenocassrpy in melons would help to elucidate the genetic mechanism of parthenocarpy not only in cucumber but also in other cucurbit crops. The purpose of this study was to clarify the inheritance of parthenocarpic ability in oriental melon cultivars and a weedy melon line, and to identify loci controlling this trait in biparental mapping populations. We determined DNA markers linked to parthenocarpic ability for marker-assisted selection and confirmed the selection accuracy.ss

## Materials and methods

We generated F_2_ progeny by selfing of F_1_s derived from crosses between the non-parthenocarpic muskmelon ‘Earl’s Favourite Harukei 3’ (EF), in the *cantalupensis–reticulatus* group, and four parthenocarpic accessions (JP82421, JP138247, JP204614, JP215901). JP82421 is a weedy melon in the *agrestis* group collected from an island in the Seto Inland Sea. JP138247, JP204614, JP215901 are old cultivars in the *conomon* or *makuwa* groups. For QTL analyses of parthenocarpic ability, we used 312 F_2_ plants derived from F_1_ of EF ♀ × JP82421 ♂ (156 in September–December 2013, 156 in March–July 2014), 291 F_2_ plants from F_1_ of EF ♀ × JP215901 ♂ (136 in September–December 2015, 155 in August–November 2018), and 307 F_2_ plants from F_1_ of JP138247 ♀ × EF ♂ (163 in February–May 2018, 144 in August–November 2018). To assess the availability of SNP markers for parthenocarpic ability, we used another two F_2_ populations: 152 plants from F_1_ of JP204614 ♀ × EF ♂ in August–November 2014, and 98 plants from F_1_ of JP138247 ♀ × EF ♂ in September–December 2017. In each crop, we also grew four to ten plants of the parental lines and their F_1_s.

### Phenotyping of parthenocarpic ability

Plants were grown on the farms of the Tsukuba Plant Innovation Research Center, University of Tsukuba (Tsukuba, Ibaraki, Japan). This site has an extratropical climate with four distinct seasons. Polyhouses were heated to keep the air temperature above 10 °C from late autumn to early spring. In each crop, seeds were sown in 6.0-or 7.5-cm-diameter plastic pots filled with moist culture soil (Nihon Hiryo Co., Ltd., Tokyo, Japan), and seedlings were raised first in an artificial climate chamber for 10 to 20 days. To minimize the effect of nonuniformity of soil conditions on plant growth and fruit development, seedlings were transplanted into 21-cm-diameter plastic pots filled with a 2:1 mixture of peat moss soil (Sakata Seed Corp., Yokohama, Japan) and culture soil in 2013–2016, and into rectangular planter boxes (30 L) filled with peat moss soil (Hokkaido Peat Moss Inc., Konosu, Japan) in 2017–2018.

At ∼25 days after transplanting, we started stigma excision of flower buds (1–3 days before flowering) at the first node of the first lateral branches above the 8th to 21st nodes of the main stem (Yoshioka et al. 2018). If most of these flowers were male, we instead used hermaphroditic (or female) flowers at the first node of the second lateral branches developed from the first node of lateral branches at the 6th to 21st nodes of the main stem. A month after the last treatment of each plant, we counted the number of enlarged fruits that were intact. To avoid erroneous assessments due to adventitious fruit development such as unexpected pollination before stigma excision or environmental factors, plants bearing two or more fruits were classified as having parthenocarpic ability.

### ddRAD-Seq analysis

Genomic DNA was isolated from individuals of parental lines and F_2_ populations with a DNeasy Plant Kit 96 (Qiagen, Hilden, Germany). Each DNA sample was diluted to a final concentration of 25 ng/μL. Library construction and sequencing analysis were performed as described by Shirasawa et al. (2016) with minor modifications. The ddRAD-Seq libraries were constructed with two combinations of restriction enzymes, *Pst*I and *Msp*I (Thermo Fisher Scientific, Waltham, MA, USA). Digested DNA was ligated to adapters with T4 DNA ligase (Takara Bio Inc., Shiga, Japan) and purified with Agencourt AMPure XP cleanup reagent (Beckman Coulter, Brea, CA, USA) to eliminate short (>300 bp) fragments. PCR amplified purified DNA with indexed primers. Amplified DNA fragments were purified with a QIAquick PCR Purification Kit (Qiagen), and fragments 300–1000 bp in length were fractionated by electrophoresis in 2.0% agarose gel (Kanto Chemical Co., Inc., Tokyo, Japan) with a MinElute Gel Extraction Kit (Qiagen). These libraries were sequenced on a HiSeq 4000 sequencer (Illumina Inc., San Diego, CA, USA) in 100-bp paired-end reads.

### Linkage map construction and QTL analysis

A genetic linkage map was constructed for each F_2_ population from the SNPs obtained by ddRAD-seq analysis in JoinMap v. 4.0 software. The SNP markers were selected by removing low-quality loci (>80% SNP progeny with missing values at the locus; mapped to Chr. 0). F_2_ individuals with >80% SNP markers were selected. Then *P*-values were calculated to test for deviations of genotype frequencies at each locus from the expected ratio of 1:2:1, and markers that showed significant distortion at the 5% level were removed. SNP markers physically mapped to each chromosome were separately used to form each linkage group. Kosambi’s mapping function was applied during the calculation of genetic map distance. QTL analysis for parthenocarpic ability was performed by nonparametric interval mapping under a single QTL model in R/qtl software (Arends et al. 2010). We calculated the 1.5-LOD support interval for the location of each QTL.

### Prediction of candidate genes, sequencing analysis, and CAPS analysis

Candidate genes within the 1.5-LOD intervals of QTLs were extracted from the Cucurbit Genomics Database (CuGenDB, http://cucurbitgenomics.org/). We designed 28 primers for sequencing two randomly selected genes related to phytohormones in the regions on Chrs. 2 and 3. Genes were amplified in 10 μL containing 2 μL of total DNA (5 ng/μL), 5 μL of 2× Ampdirect Plus (Shimadzu, Kyoto, Japan), 0.1 μL of Blend Taq -Plus-(2.5 U/μL) (Toyobo, Osaka, Japan), and 10 μM each forward and reverse primer. The thermal profile consisted of 95 □ for 2 min; 35 cycles at 95 □ for 30 s, 55 □ for 30 s, and 72 □ for 1 min; and a final extension of 72 □ for 5 min. For sequencing analysis of the gene on Chr. 2, the PCR products were Sanger-sequenced by Fasmac Co., Ltd. (Kanagawa, Japan). For CAPS analysis of the gene on Chr. 3, according to sequence polymorphisms with parental lines, we used a previously developed CAPS marker (Nashiki et al. 2023), permitting digestion with *Alu*□ from the 5′ end of both EF alleles but not from parthenocarpic accessions. Finally, the digested PCR products were electrophoresed in 2.0% agarose gel in 1× TAE buffer.

## Results

### Segregation of parthenocarpic ability

The non-parthenocarpic ‘Earl’s Favourite Harukei 3’ (EF) consistently bore no fruits, whereas all plants of parthenocarpic accessions showed parthenocarpic ability except for one plant of JP138247 in September– December 2017 (Tables S1, S2). The F_1_ plants of EF × parthenocarpic accessions also showed parthenocarpic ability except for one plant of JP138247 ♀ × EF ♂ and two plants of JP204614 ♀ × EF ♂. Although the plants of parental lines and their F_1_s bore bisexual flowers at the first nodes of most first lateral branches, several plants of the F_2_ populations bore many male flowers at the first nodes of both first and second lateral branches. Such plants, without enough bisexual flowers, were excluded from subsequent analyses. In F_2_ of EF ♀ × JP82421 ♂, 154 plants were phenotyped in 2013 (72 parthenocarpic ability), and 150 in 2014 (48) (Fig. 1; Tables S1, S2). In F_2_ of EF ♀ × JP215901 ♂, 130 plants were phenotyped in 2015 (41 parthenocarpic ability), and 130 in 2018 (59). In F_2_ of JP138247 ♀ × EF ♂, 83 plants were phenotyped in 2017 (39 parthenocarpic ability), 151 in February– May 2018 (104), and 133 in August–November 2018 (70). In F_2_ of JP204614 ♀ × EF ♂, 148 plants were phenotyped in 2014 (55 parthenocarpic ability). Chi-squared analysis indicated that all of these ratios differed significantly from 3:1 (single gene) except for F_2_ of EF ♀ × JP215901 ♂ in 2015 and F_2_ of JP138247 ♀ × EF ♂ in February–May 2018.

**Fig. 1.**
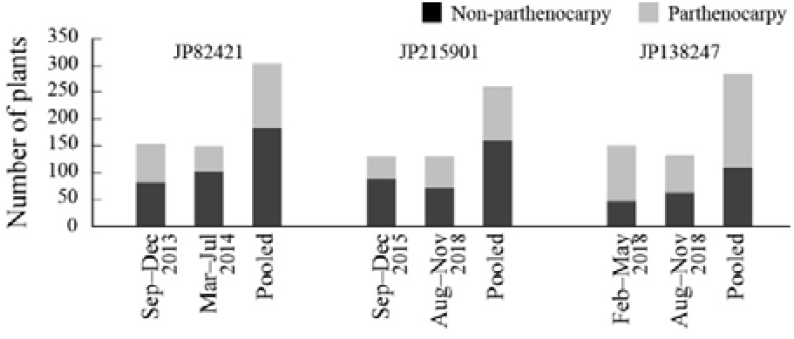
Phenotypes of three F_2_ populations derived from a cross between a non-parthenocarpic cultivar and three parthenocarpic cultivars/lines.

### Linkage map and QTL analysis

We obtained 355□358□197 reads from plants in F_2_ of EF ♀ × JP82421 ♂ (1677 SNPs); 348□703□237 reads in F_2_ of EF ♀ × JP215901 ♂ (3079 SNPs); and 163□841□414 reads in F_2_ of JP138247 ♀ × EF ♂ (2883 SNPs).

After filtering, linkage maps were constructed with SNP markers in F_2_ of EF ♀ × JP82421 ♂ (1331 markers), F_2_ of EF ♀ × JP215901 ♂ (1161 markers), and F_2_ of JP138247 ♀ × EF ♂ (1163 markers). All maps contained 12 linkage groups spanning 1305.9, 1463.7, and 1280.0 cM, with an average marker distance of 1.0, 1.3, and 1.1 cM, respectively. In F_2_ of EF ♀ × JP82421 ♂, we detected two QTLs for parthenocarpic ability on Chrs. 2 and 3 in both crops (Fig. 2; Table 1). QTL analysis using pooled data detected an additional QTL on Chr. 7. The peak LOD of the QTL on Chr. 3 was slightly lower than that of the QTL on Chr. 2 in 2014, but higher in 2013 and pooled data. In F_2_ of EF ♀ × JP215901 ♂, we detected only one QTL on Chr. 3 in both crops. QTL analysis using pooled data detected an additional QTL on Chr. 11. The peak LOD of the QTL on Chr. 3 was much higher than that of the QTL on Chr. 11 in the pooled data. In F_2_ of JP138247 ♀ × EF ♂, we detected two QTLs on Chrs. 2 and 3 in both crops and in the pooled data. The peak LODs of the QTLs on Chr. 3 were higher than those of the QTLs on Chr. 2 in both crops and in the pooled data. Comparison of the physical positions of the QTLs indicated that those on Chr. 2 (*par2.1*) detected in two populations were identical and those on Chr. 3 (*par3.1*) detected in all three populations were identical (Table 1). The parthenocarpic parent allele at both QTLs enhanced parthenocarpic ability. Since *par2.1* and *par3.1* were stably detected in different crops in multiple populations, we focused on both in subsequent analyses. Parthenocarpic ability (ratio of non-parthenocarpic to parthenocarpic plants) was well explained by alleles at both QTLs (genotyped by nearest SNPs to LOD peak in each QTL region) in F_2_s of EF ♀ × JP82421 ♂ and of JP138247 ♀ × EF ♂ (Fig. S1), but by only the QTL on Chr. 3 in F_2_ of EF ♀ × JP215901 ♂.

**Fig. 2.**
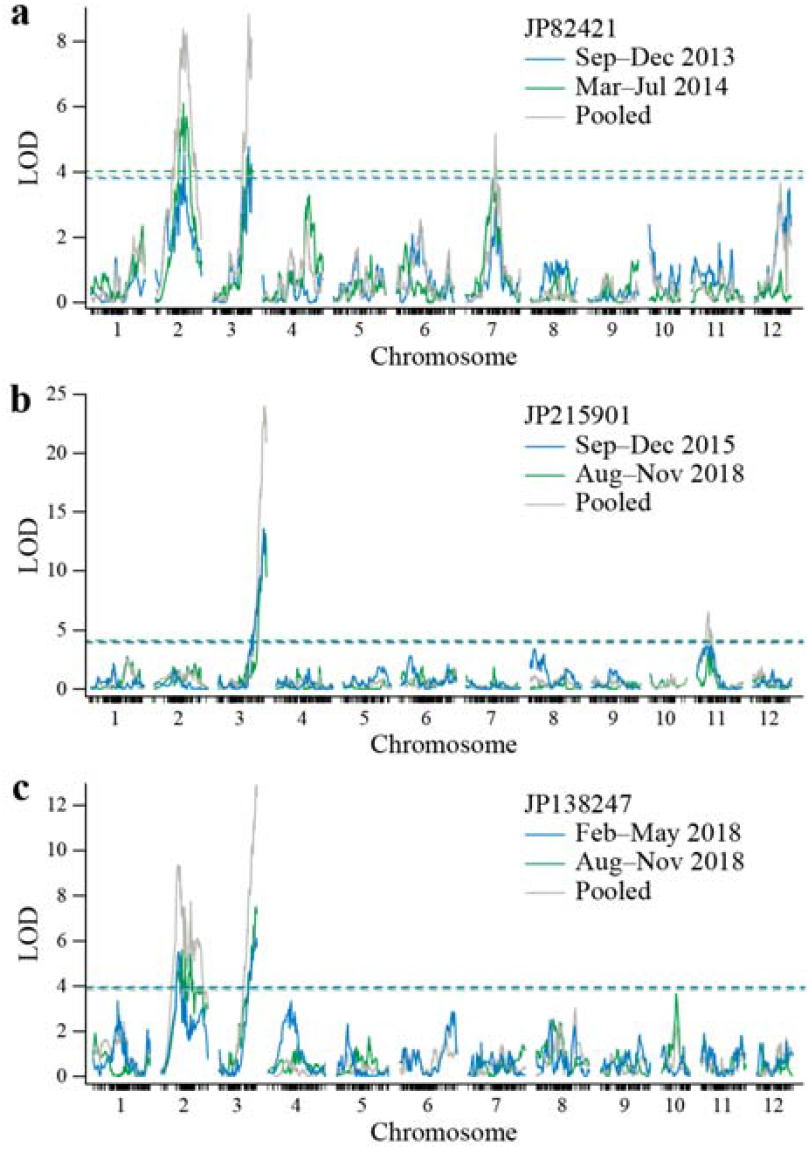
Positions and LOD scores of QTLs associated with parthenocarpic ability in F_2_ populations of (**a**) ‘Earl’s Favourite Harukei 3’ (EF) ♀ × JP82421 ♂, (**b**) EF ♀ × JP215901 ♂, and (**c**) JP138247 ♀ × EF ♂. Horizontal lines show LOD thresholds at *P* = 0.05.

**Table 1.** QTLs for parthenocarpic ability detected by non-parametric single-interval mapping in F_2_ populations derived from crosses between parthenocarpic accessions and a non-parthenocarpic cultivar.

| F<sub>2</sub> population and crop | Chr. | Position (cM) | LOD peak | Estimated range within 'DHL92 v. 3.5.1' genome |
| --- | --- | --- | --- | --- |
| F<sub>2</sub> 'Earl's Favourite Harukei 3' ♀ × JP82421 ♂ |  |  |  |  |
| September–December 2013 | 2 | 64.1 | 4.5 | 21617006–21594579 |
|  | 3 | 81.7 | 4.8 | 28360965–28546200 |
| March–July 2014 | 2 | 62.2 | 6.1 | 15371188–23307877 |
|  | 3 | 78.6 | 4.5 | 27700246–29130876 |
| Pooled | 2 | 62.2 | 8.4 | 3048035–24438959 |
|  | 3 | 81.7 | 8.8 | 26506946–28546200 |
|  | 7 | 67.9 | 5.2 | 5184527–6074771 |
| F<sub>2</sub> 'Earl's Favourite Harukei 3' ♀ × JP215901 ♂ |  |  |  |  |
| September–December 2015 | 3 | 113 | 13.6 | 26047561–28658287 |
| August–November 2018 | 3 | 117 | 13.1 | 27234697–28658287 |
| Pooled | 3 | 115 | 24.0 | 26047561–28658287 |
|  | 11 | 29 | 6.6 | 5642934–13474946 |
| F<sub>2</sub> JP138247 ♀ × 'Earl's Favourite Harukei 3' ♂ |  |  |  |  |
| February–May 2018 | 2 | 39 | 5.5 | 2292091–2580477 |
|  | 3 | 82 | 6.1 | 25974023–28621101 |
| August–November 2018 | 2 | 46.5 | 5.6 | 2292091–24732179 |
|  | 3 | 82 | 7.5 | 26814676–28621101 |
| Pooled | 2 | 37 | 9.3 | 1987986–24732179 |
|  | 3 | 82 | 12.9 | 25320477–28621101 |

### Prediction of candidate genes, sequencing analysis, and CAPS analysis

Using the melon genome ‘DHL92’ v. 3.5.1 annotation database, we annotated genes within the 1.5-LOD intervals of the QTLs at *par2.1* (92 genes, Table S3) and *par3.1* (38 genes, Table S4). Both regions contain several phytohormone-related genes. We randomly selected one gene in each region for developing SNP markers for marker-assisted selection of parthenocarpic ability. We selected MELO3C017414 (encoding an auxin efflux carrier component) at *par2.1*, and sequenced it in EF and the parthenocarpic accessions. We found two SNPs in its coding region between EF and the parthenocarpic accessions, but none among the latter. We also selected and sequenced MELO3C010779 (encoding 1-aminocyclopropane-1-carboxylate synthase) at *par3.1*, well known as *CmACS11*, which catalyzes the rate-limiting step in ethylene biosynthesis (Boualem et al. 2015), and detected polymorphisms between EF and the parthenocarpic accessions. We found eight SNPs, including three non-synonymous substitutions, in the coding region of *CmACS11* between EF and the parthenocarpic accessions, but none among the latter.

### Reconfirmation of SNP markers for selection of parthenocarpic ability

We used a SNP at 132 bp from the start codon in MELO3C017414 for genotyping *par2.1* and one at 1050 bp from the start codon in *CmACS11* for genotyping *par3.1* in two additional populations: F_2_ of JP204614 ♀ × EF ♂, and F_2_ of JP138247 ♀ × EF ♂. Genotyping data were obtained from each F_2_ plant by sequencing for *par2.1* and by a previously developed CAPS marker for *par3.1* (Nashiki et al. 2023). The ratios of non-parthenocarpic to parthenocarpic plants in all two populations were well explained by the combination of these two markers (Fig. 3).

**Fig. 3.**
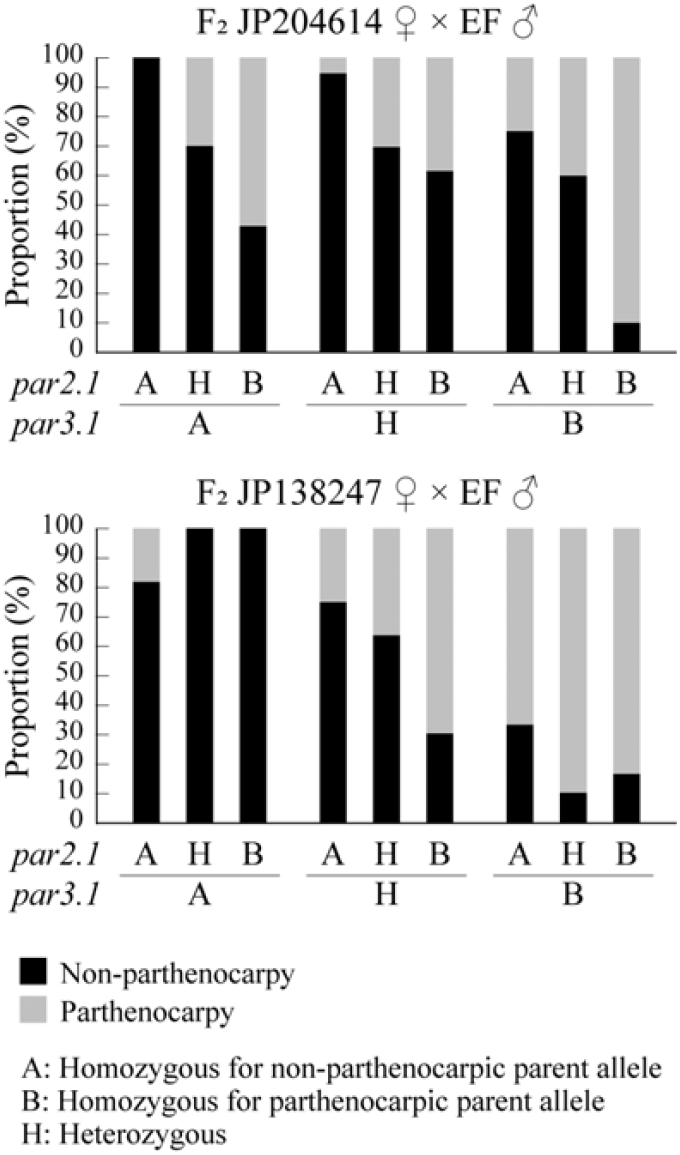
Relationships of parthenocarpic ability with genotype at two QTLs on chromosomes 2 (*par2.1*) and 3 (*par3.1*) in three F_2_ populations derived from a cross between non-parthenocarpic ‘Earl’s Favourite Harukei 3’ (EF) and two parthenocarpic accessions (JP204614, JP138247). Genotype of each F_2_ plant was determined by SNPs in MELO3C017414 for *par2.1* and in *CmACS11* for *par3.1*.

## Discussion

Natural (genetic) parthenocarpy observed in East Asian melon accessions seems to be controlled by two major QTLs on Chrs. 2 (*par2.1*) and 3 (*par3.1*). The few mutations in the sequences of the randomly selected genes in the two QTL regions among accessions suggest a high likelihood that the parthenocarpic genes within the accessions examined here are identical. Screening for parthenocarpic ability by Yoshioka et al. (2018) showed several accessions derived from China, Korea, and Japan to have strong parthenocarpy, in addition to the four accessions that we used, suggesting that the genes on Chrs. 2 and 3 may be common in East Asian melons. According to a recent analysis of diverse melon accessions from diverse regions, two or three independent domestication events occurred in India and Africa (Endl et al. 2018; Zhao et al. 2019; Wang et al. 2021). East Asian melon is likely derived from Indian *agrestis* melons (Wang et al. 2021), and was introduced via China into Japan and Korea. Although it is unclear when and where in the processes of domestication and spread it obtained parthenocarpic ability, the genes on Chrs. 2 and 3 have been conserved for a long time in East Asian melon. Parthenocarpy assures yields because fruit develops even under unfavorable conditions, and thus parthenocarpic genes may have been unconsciously selected.

We detected *par2.1* and *par3.1* in multiple crops, with large effects. Phenotypic segregation in the F_2_ populations was well explained by their combination in the F_2_s of EF ♀ × JP82421 ♂ and of JP138247 ♀ × EF ♂ (Fig. S1), and by *par3.1* alone in the F_2_ of EF ♀ × JP215901 ♂ (Fig. S1). In the former populations, all plants homozygous for parthenocarpic parent alleles at both *par2.1* and *par3.1* showed high parthenocarpic ability. In the latter population, most plants that were homozygous for the parthenocarpic parent allele of *par3.1* showed high parthenocarpic ability. The difference in the effects of *par2.1* and *par3.1* may be due to differences in genetic background among accessions, but further experiments will be required to test this hypothesis. In cucumber, many QTLs have been detected in analyses using genetically distinct parthenocarpic cucumbers (Sun et al. 2006; Lietzow et al. 2016; Wu et al. 2016). The low number of QTLs detected here may be due to differences in the methods of evaluation rather than in genetic mechanisms between species: that is, the studies of cucumber evaluated parthenocarpic ability quantitatively by parthenocarpic fruit yield or the proportion of parthenocarpic fruit set, whereas we used binary scores of parthenocarpic ability in our QTL analyses, and neglected quantitative variation in parthenocarpic ability. Therefore, if we evaluated parthenocarpic ability of melon using the same method as in the cucumber studies, we might detect new loci related to quantitative variation in parthenocarpy. Moreover, parthenocarpic ability was segregated even among F_2_ plants considered to be of the same genotype at both *par2.1* and *par3.1*. This suggests the possibility of other minor-effect loci, such as QTLs on Chr. 7 and 11 detected only by QTL analyses using pooled data.

On the other hand, the influence of environment cannot be ignored. It is well known that parthenocarpy in cucumber and squash is influenced by the environment (Robinson and Reiners 1999; Gou et al. 2022). Yoshioka et al. (2018) reported that parthenocarpic melon accessions had stable parthenocarpic ability throughout the year, but the size and number of parthenocarpic fruits differed among seasons, indicating the importance of environmental factors for parthenocarpic fruit development in melon. Here, all plants of parthenocarpic accessions, except for one of JP138247, consistently showed strong parthenocarpic ability in different crops, whereas those of the non-parthenocarpic cultivar did not. However, the ratios of non-parthenocarpic to parthenocarpic F_2_ plant numbers differed significantly between crops in each population. We did not collect environmental data in the polyhouses, and thereby we cannot clearly state which environmental factors differed among crops, but it is possible that the influence of some factors affected the expression of parthenocarpy. Moreover, segregation of parthenocarpic ability among F_2_ plants with the same genotype at the parthenocarpy QTLs suggests that subtle differences in the environment around each plant would affect fruit set.

Several phytohormone-related genes occur in the 1.5-LOD support intervals of each QTL (Tables S3, S4), and more occur outside of these intervals. Several phytohormones have been shown to induce parthenocarpic fruit development in a number of species (Joldersma and Liu 2018). In horticultural crops, auxins, gibberellins, and cytokinins are primary factors in initiating fruit set, and other hormones such as ethylene, brassinosteroids, and melatonin also have significant effects on parthenocarpic fruit formation (Sharif et al. 2022). We randomly selected one phytohormone-related gene within each QTL interval for the development of DNA markers. It will be necessary to conduct further experiments to clarify the relationship between expression of other phytohormone-related genes and parthenocarpic ability.

As in Yoshioka et al. (2018), most F_1_ plants of crosses between the non-parthenocarpic cultivar and the parthenocarpic accessions did not show strong parthenocarpic ability, indicating that the ability in East Asian melon is recessive. Therefore, to confer parthenocarpy on an F_1_ cultivar, both of its parents must have parthenocarpic ability. This means that more populations need to be treated in breeding programs. In addition, a considerable number of F_2_ plants without parthenocarpic alleles at either *par2.1* or *par3.1* showed parthenocarpic ability, indicating a high possibility that phenotypic selection missed those alleles. Therefore, the use of DNA markers tightly linked to requisite parthenocarpic genes is an important requirement for the efficient breeding of new cultivars. On this point, we showed that the SNP markers used here or other SNP markers in the QTL region are sufficient, though not perfect, to select plants with parthenocarpic genes at both *par2.1* and *par3.1*. Elucidation of the network of hormones and these two genes in the control of parthenocarpic ability in melon may provide new insights into the genetic mechanism of parthenocarpy in cucurbits.

## Supporting information

Online resource 1

## Acknowledgments

This work was funded by JSPS KAKENHI Grant Numbers 23780034 and 26712004 to YY. We are grateful to N. Nishi, H. Matsumura, T. Iwaya, M. Ishida and K. Moriyama of the University of Tsukuba for technical assistance.

## Data availability

Datasets generated and analyzed during this study are available from the corresponding author on reasonable request.

