## Supplementary material for "Identification of major quantitative trait loci for parthenocarpic ability in East Asian melon": Online resource 1

### Supplementary materials

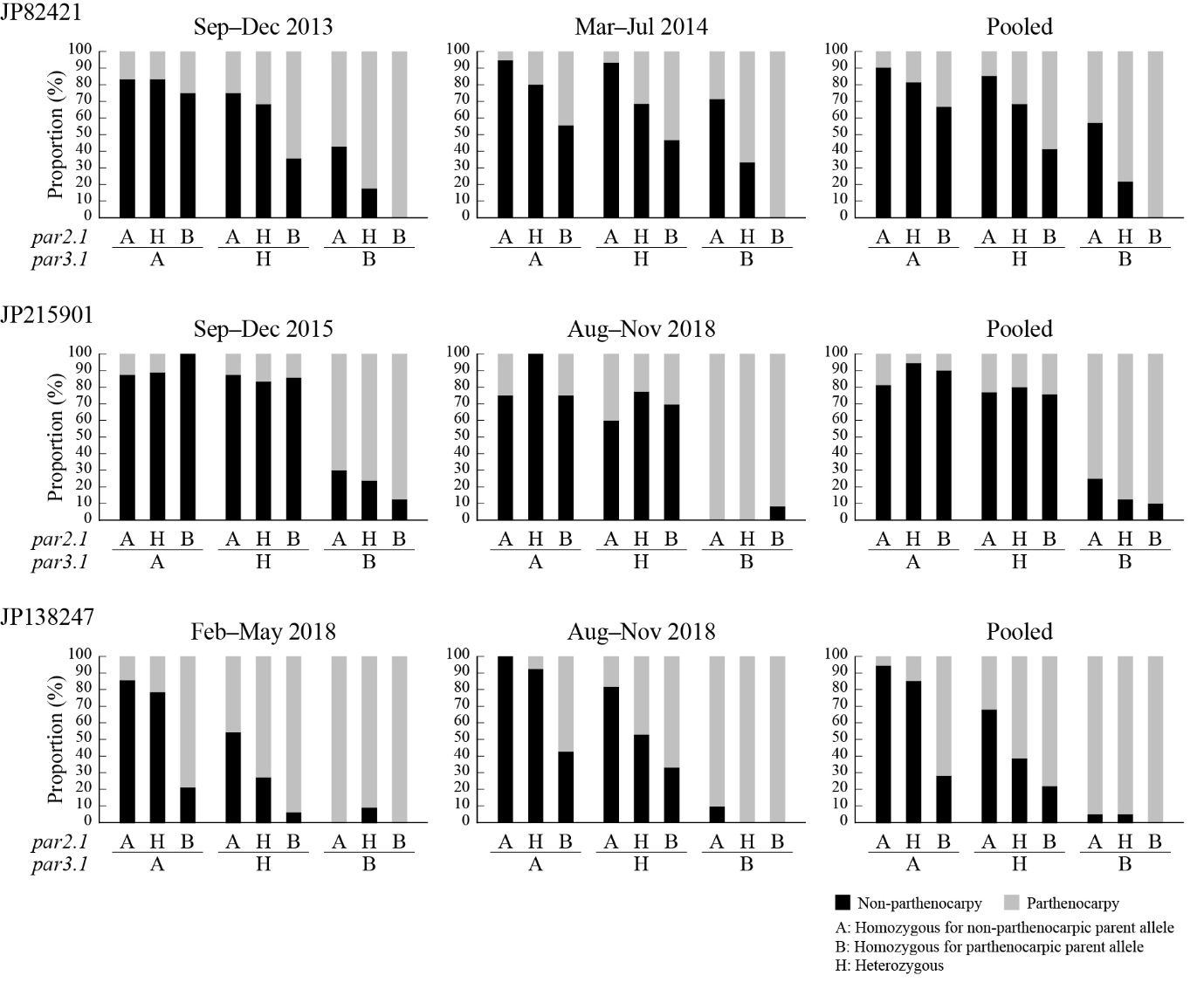

**Fig. S1** Relationships of parthenocarpic ability (ratios of non-parthenocarpic to parthenocarpic plants) with genotype at two QTLs on chromosomes 2 (*par2.1*) and 3 (*par3.1*) in three F_2_ populations used in QTL analyses. Genotypes were determined by the nearest SNPs to the LOD peak in each QTL region.

**Table S1** Parthenocarpic ability of parental lines and their F_1_ and F_2_ progeny by QTL analysis in two crops of each of three populations.

| Parental lines/progeny | No. of plants^1^ | Non-parthenocarpy | Parthenocarpy |
| --- | --- | --- | --- |
| *September–December 2013* |  |  |  |
| ‘Earl’s Favourite Harukei 3’ (EF) | 6 | 6 | 0 |
| JP82421 | 6 | 0 | 6 |
| F_1_ EF ♀ × JP82421 ♂ | 6 | 6 | 0 |
| F_2_ EF ♀ × JP82421 ♂ | 154 | 82 | 72 |
| *March–July 2014* |  |  |  |
| ‘Earl’s Favourite Harukei 3’ (EF) | 8 | 8 | 0 |
| JP82421 | 8 | 0 | 8 |
| F_1_ EF ♀ × JP82421 ♂ | 8 | 8 | 0 |
| F_2_ EF ♀ × JP82421 ♂ | 150 | 102 | 48 |
| *September–December 2015* |  |  |  |
| ‘Earl’s Favourite Harukei 3’ (EF) | 8 | 8 | 0 |
| JP215901 | 8 | 0 | 8 |
| F_1_ EF ♀ × JP215901 ♂ | 8 | 8 | 0 |
| F_1_ JP215901 ♀ × EF ♂ | 8 | 8 | 0 |
| F_2_ EF ♀ × JP215901 ♂ | 130 | 89 | 41 |
| *August–November 2018* |  |  |  |
| ‘Earl’s Favourite Harukei 3’ (EF) | 5 | 5 | 0 |
| JP215901 | 5 | 0 | 5 |
| F_1_ EF ♀ × JP215901 ♂ | 5 | 5 | 0 |
| F_1_ JP215901 ♀ × EF ♂ | 5 | 5 | 0 |
| F_2_ EF ♀ × JP215901 ♂ | 130 | 71 | 59 |
| *February–May 2018* |  |  |  |
| ‘Earl’s Favourite Harukei 3’ (EF) | 5 | 5 | 0 |
| JP138247 | 5 | 0 | 5 |
| F_1_ EF ♀ × JP138247 ♂ | 5 | 5 | 0 |
| F_1_ JP138247 ♀ × EF ♂ | 5 | 4 | 1 |
| F_2_ JP138247 ♀ × EF ♂ | 151 | 47 | 104 |
| *August–November 2018* |  |  |  |
| ‘Earl’s Favourite Harukei 3’ (EF) | 5 | 5 | 0 |
| JP138247 | 5 | 0 | 5 |
| F_1_ EF ♀ × JP138247 ♂ | 5 | 5 | 0 |
| F_1_ JP138247 ♀ × EF ♂ | 5 | 5 | 0 |
| F_2_ JP138247 ♀ × EF ♂ | 133 | 63 | 70 |
| ^1^ Number of plants that could be evaluated. | |  |  |

**Table S2** Parthenocarpic ability of parental lines and their F_1_ and F_2_ progeny for confirmation of marker selection accuracy.

| Parental lines/progeny | No. of plants^1^ | Non-parthenocarpy | Parthenocarpy |
| --- | --- | --- | --- |
| *August–November 2014* |  |  |  |
| ‘Earl’s Favourite Harukei 3’ (EF) | 7 | 7 | 0 |
| JP204614 | 7 | 0 | 7 |
| F_1_ EF ♀ × JP204614 ♂ | 7 | 7 | 0 |
| F_1_ JP204614 ♀ × EF ♂ | 7 | 5 | 2 |
| F_2_ JP204614 ♀ × EF ♂ | 148 | 93 | 55 |
| *September–December 2017* |  |  |  |
| ‘Earl’s Favourite Harukei 3’ (EF) | 3 | 3 | 0 |
| JP138247 | 3 | 1 | 2 |
| F_1_ EF ♀ × JP138247 ♂ | 4 | 4 | 0 |
| F_1_ JP138247 ♀ × EF ♂ | 4 | 4 | 0 |
| F_2_ JP138247 ♀ × EF ♂ | 83 | 44 | 39 |
| ^1^ Number of plants that could be evaluated. | |  |  |

**Table S3** List of genes within the 1.5-LOD support interval of the QTL for parthenocarpic ability on chromosome 2 in three populations.

| Gene ID | Description (DHL v3.5.1) |
| --- | --- |
| MELO3C017499 | 1-deoxy-D-xylulose-5-phosphate synthase |
| MELO3C017498 | Alpha/beta-hydrolase superfamily protein |
| MELO3C017497 | Cysteine-rich venom protein |
| MELO3C017496 | Cysteine-rich venom protein |
| MELO3C017495 | UDP-glucosyltransferase, putative |
| MELO3C017494 | Myb/SANT-like DNA-binding domain protein |
| MELO3C017493 | Pectin lyase-like superfamily protein |
| MELO3C017492 | Luciferin 4-monooxygenase |
| MELO3C017491 | Myb transcription factor |
| MELO3C017490 | Putative aminotransferase |
| MELO3C017489 | F-box family protein, putative |
| MELO3C017488 | Guanine nucleotide-binding protein subunit beta |
| MELO3C017487 | F-box protein PP2-B1 |
| MELO3C017486 | F-box protein PP2-B1 |
| MELO3C017485 | F-box protein PP2-B1 |
| MELO3C017484 | F-box protein PP2-B1 |
| MELO3C017483 | Calmodulin-binding family protein |
| MELO3C017482 | Xyloglucan endotransglucosylase/hydrolase |
| MELO3C017481 | Xyloglucan endotransglucosylase/hydrolase |
| MELO3C017480 | Xyloglucan endotransglucosylase/hydrolase |
| MELO3C017479 | Xyloglucan endotransglucosylase/hydrolase |
| MELO3C017478 | Xyloglucan endotransglucosylase/hydrolase |
| MELO3C017477 | Acyl-[acyl-carrier-protein] desaturase 7, chloroplastic |
| MELO3C017476 | Xyloglucan endotransglucosylase/hydrolase |
| MELO3C017475 | Nicotianamine synthase |
| MELO3C017474 | zinc finger protein 2 |
| MELO3C017473 | 4-coumarate:CoA ligase 1 |
| MELO3C017472 | Response regulator |
| MELO3C017471 | Mitochondrial outer membrane porin 4-like protein |
| MELO3C017470 | Putative beta-1,3-galactosyltransferase 16 |
| MELO3C017469 | Voltage-dependent L-type calcium channel subunit alpha-1C |
| MELO3C017467 | Syntaxin-61-like protein |
| MELO3C017468 | NAD-dependent glycerol-3-phosphate dehydrogenase family protein |
| MELO3C017466 | Small, acid-soluble spore protein Tlp |
| MELO3C017465 | FCH domain only protein 1 |
| MELO3C017464 | Activating signal cointegrator 1 complex subunit 1 |
| MELO3C017463 | Protein kinase superfamily protein with octicosapeptide/Phox/Bem1p domain |
| MELO3C017462 | Haloacid dehalogenase-like hydrolase |
| MELO3C017461 | NADPH--cytochrome P450 reductase |
| MELO3C017460 | Chromosome 3B, genomic scaffold, cultivar Chinese Spring |
| MELO3C017459 | Protein of unknown function (DUF688) |
| MELO3C017458 | Replication protein A 32 kDa subunit-like protein |
| MELO3C017457 | AP2-like ethylene-responsive transcription factor |
| MELO3C017456 | ABC transporter, ATP-binding protein |
| MELO3C017455 | Protein VERNALIZATION INSENSITIVE 3 |
| MELO3C017454 | U4/U6.U5 small nuclear ribonucleoprotein 27 kDa protein |
| MELO3C017453 | Putative receptor-like kinase |
| MELO3C017452 | Transducin/WD-like repeat-protein |
| MELO3C017451 | Hydroxyproline-rich glycoprotein family protein |
| MELO3C017450 | H+-ATPase family protein |
| MELO3C017449 | A-kinase anchor protein 9, putative isoform 2 |
| MELO3C017448 | Putative glucose-6-phosphate 1-epimerase |
| MELO3C017447 | Villin-4-like protein |
| MELO3C017446 | Cysteine/Histidine-rich C1 domain family protein |
| MELO3C017445 | Hepatocyte growth factor-regulated tyrosine kinase substrate |
| MELO3C017444 | RNA helicase family protein |
| MELO3C017443 | Cyclic pyranopterin monophosphate synthase accessory protein |
| MELO3C017442 | Calcium-dependent lipid-binding (CaLB domain) plant phosphoribosyltransferase family protein |
| MELO3C017441 | Aerobic glycerol-3-phosphate dehydrogenase |
| MELO3C017440 | U3 small nucleolar RNA-associated protein 25 |
| MELO3C017439 | Metaxin-related family protein |
| MELO3C017438 | Putative copper-transporting ATPase 3 |
| MELO3C017437 | Core-2/I-branching beta-1,6-N-acetylglucosaminyltransferase family protein |
| MELO3C017436 | Eukaryotic aspartyl protease family protein, putative |
| MELO3C017435 | Eukaryotic aspartyl protease family protein, putative |
| MELO3C017434 | Oxysterol-binding protein-related protein 4C |
| MELO3C017433 | Transmembrane protein, putative |
| MELO3C017432 | Serine/arginine repetitive matrix protein 2, putative isoform 1 |
| MELO3C017431 | Pectinesterase |
| MELO3C017430 | Fiber protein Fb15 |
| MELO3C017429 | Pathogenesis-related homeodomain protein |
| MELO3C017428 | RING/FYVE/PHD zinc finger-containing protein |
| MELO3C017427 | tRNA-specific adenosine deaminase 2 |
| MELO3C017426 | Macrophage migration inhibitory factor like |
| MELO3C017425 | Proline-rich PRCC |
| MELO3C017424 | Transcription factor |
| MELO3C017423 | Ubiquitin-fold modifier-conjugating enzyme 1 |
| MELO3C017422 | Purple acid phosphatase 28 |
| MELO3C017421 | Putative inactive purple acid phosphatase 28 |
| MELO3C017420 | ATP binding, related |
| MELO3C017419 | DNA ligase |
| MELO3C017418 | cysteine-rich RLK (RECEPTOR-like protein kinase) 29 |
| MELO3C017417 | Transmembrane-like protein |
| MELO3C017416 | Cyclin-dependent protein kinase inhibitor Siamese |
| MELO3C017415 | WRKY transcription factor 11 |
| MELO3C017414 | Putative auxin efflux carrier protein |
| MELO3C017413 | Oligosaccharyltransferase complex subunit OSTC-like protein |
| MELO3C017412 | Leucine-rich repeat (LRR) family protein |
| MELO3C017411 | Glutaredoxin |
| MELO3C017410 | Glutaredoxin |
| MELO3C017409 | Bifunctional lysine-specific demethylase and histidyl-hydroxylase NO66 |
| MELO3C017408 | Short transient receptor potential channel 4-associated protein |

**Table S4** List of genes within the 1.5-LOD support interval of the QTL for parthenocarpic ability on chromosome 3 in three populations.

| Gene ID | Description (DHL v3.5.1) |
| --- | --- |
| MELO3C010816 | Protein kinase 2 |
| MELO3C010815 | Peroxisomal membrane MPV17/PMP22-like protein |
| MELO3C010814 | Family of Uncharacterized protein function (DUF566), putative isoform 1 |
| MELO3C010813 | Zinc-finger protein 1 |
| MELO3C010812 | Adenine phosphoribosyltransferase 1 |
| MELO3C010811 | Transcription factor, putative |
| MELO3C010810 | Pimeloyl-[acyl-carrier protein] methyl ester esterase |
| MELO3C010809 | U-box domain-containing 4-like protein |
| MELO3C010808 | Subtilisin-like protease |
| MELO3C010807 | F-box family protein |
| MELO3C010806 | WD40 repeat-like protein |
| MELO3C010805 | GTPase family protein |
| MELO3C010804 | Tetratricopeptide repeat protein 7B |
| MELO3C010803 | Phosphatidylinositol 4-kinase beta |
| MELO3C010802 | Protein ZCW7 |
| MELO3C010801 | LIM domain protein LIM-2 |
| MELO3C010800 | F-box protein |
| MELO3C010799 | Exocyst subunit exo70 family protein G1 |
| MELO3C010798 | NADH dehydrogenase iron-sulfur protein 4 |
| MELO3C010797 | NACHT, LRR and PYD domains-containing protein 5 |
| MELO3C010796 | Coffea canephora DH200=94 genomic scaffold, scaffold_6 |
| MELO3C010795 | Calcium-transporting ATPase |
| MELO3C010794 | Calcium-transporting ATPase |
| MELO3C010793 | Acetyl-CoA decarbonylase/synthase complex subunit beta 1 |
| MELO3C010792 | DHFS-FPGS homolog D |
| MELO3C010791 | U-box domain-containing protein 30 |
| MELO3C010790 | V-type proton ATPase subunit G3 |
| MELO3C010789 | CW7 protein, putative |
| MELO3C010788 | BZIP transcription factor family protein |
| MELO3C010787 | Kinesin-like protein |
| MELO3C010786 | Growth-regulating factor 1 |
| MELO3C010785 | Glutathione transport system permease protein GsiD |
| MELO3C010784 | AP2-like ethylene-responsive transcription factor |
| MELO3C010783 | Transcription factor |
| MELO3C010782 | Cellulose synthase-like C1-1, glycosyltransferase family 2 protein |
| MELO3C010781 | Squalene monooxygenase |
| MELO3C010780 | Chaperone protein DnaJ |
| MELO3C010779 | 1-aminocyclopropane-1-carboxylate synthase |

**Table S5** Primers for sequencing and CAPS analysis used in this study.

| Primer | Sequence (5'-3') | Length(bp) | Amplicon (bp) | Notes |
| --- | --- | --- | --- | --- |
| ACS11_1F | AGCGAATAAAGTCACAAAACTG | 22 | 779 | Sequencing for CmACS11 |
| ACS11_1R | TCTCAATCCTAAAGCATCTGG | 21 |  |  |
| ACS11_2F | GGAGTGGTTTCTTTTGTGCT | 20 | 678 | Sequencing for CmACS11 |
| ACS11_2R | TGTTTGAACTCGAACAATGG | 20 |  |  |
| ACS11_3F | ATATCTTGTCTTGCCGATCC | 20 | 846 | Sequencing for CmACS11 |
| ACS11_3R | AAAGCTAGACATTTTGGTAGCC | 22 |  |  |
| ACS11_4F | GCTTTTGTTTAGTTTCTTCTCAAA | 24 | 770 | Sequencing for CmACS11 |
| ACS11_4R | AGCAAAAAGATGTGCATGAT | 20 |  |  |
| ACS11_5F | AACTGTCATGGACCCGAACC | 20 | 518 | Sequencing for CmACS11 |
| ACS11_5R | TGGCCACCTTGAAAGTGTGT | 20 |  |  |
| CmACS11_3F | ATATCTTGTCTTGCCGATCC | 20 | 498 | CAPS marker for CmACS11 |
| CAPS_ACS11_1R | TGGATTCGGTCCCAAATTGGG | 21 |  |  |
| AUX_1F | TCATCGAGTTGAGTGGGTTAGA | 22 | 588 | Sequencing for MELO3C017414 |
| AUX_1R | AACGAGAGTGTTGGGGAGAG | 20 |  |  |
| AUX_2F | TCCCTACGCCATGAACTACC | 20 | 605 | Sequencing for MELO3C017414 |
| AUX_2R | TGGCATAAAAATCCGTTTGG | 20 |  |  |
| AUX_3F | TCTGCAGCTTCTTCTATGGTTTC | 23 | 563 | Sequencing for MELO3C017414 |
| AUX_3R | GCTGAAAACATGGGGTTTG | 19 |  |  |
| AUX_4F | AACGCCAAGAACGTCAAACT | 20 | 650 | Sequencing for MELO3C017414 |
| AUX_4R | TGACACTTGCAGGAGGCATA | 20 |  |  |
| AUX_5F | CAGGGCTGCAAGAAGTGATA | 20 | 520 | Sequencing for MELO3C017414 |
| AUX_5R | TTGGTCGAGCTACGAGTTCA | 20 |  |  |
| AUX_6F | TGCAAACTATGAACCTATAGGAAA | 24 | 506 | Sequencing for MELO3C017414 |
| AUX_6R | ATGGAAGTCGCAGCCATAAC | 20 |  |  |
| AUX_7F | AATCAAATCGACCCCGTTG | 19 | 614 | Sequencing for MELO3C017414 |
| AUX_7R | CCTGCACAAATGTACACGAAA | 21 |  |  |
| AUX_8F | TCCTTAATTTCAATTAGCCATCA | 23 | 602 | Sequencing for MELO3C017414 |
| AUX_8R | GCTCCTTCCCCCTTTGATTA | 20 |  |  |
